# Genomic, Transcriptomic, and Regulomic Analyses Do Not Support Profound Autism as a Distinct Biological Category

**DOI:** 10.64898/2026.06.01.729392

**Authors:** Tara Eicher, Ari Ne’eman, John Quackenbush

## Abstract

The Lancet Commission on the Future of Care and Clinical Research in Autism proposed the construct of “profound autism” as a recognizable subtype of autism. Supporters argue that this classification is necessary to ensure that autistic persons with severe impairment receive appropriate research attention and policy support, whereas critics contend that the construct lacks scientific validity and may reflect social or political considerations more than biological distinction. To inform this debate, we evaluate whether the proposed “profound autism” category represents a distinct genetic phenotype using multiple molecular data types collected in a large cohort. Across genomic, transcriptomic, and regulatory analyses, we find no evidence supporting “profound autism” as a biologically distinct phenotypic group. Instead, differences emerge primarily in inferred gene regulatory networks distinguishing nonspeaking from speaking autistic children, suggesting potential regulatory mechanisms contributing to speech ability. These findings suggest that future research into severe impairment may be more productive if focused on specific traits—such as speech impairment—rather than attempting to define a distinct biological subtype within the multidimensional phenomenon of autism.

## Introduction

In 2022, the *Lancet Commission on the Future of Care and Clinical Research in Autism* proposed adoption of the term “profound autism” to describe autistic children and adults who require 24- hour access to care, cannot be safely left alone, and are unable to meet basic adaptive needs independently. To operationalize this construct, the Commission defined “profound autism” as persons with either significant intellectual disability (IQ below 50), very limited language ability (for example, limited ability to communicate to a stranger using complete sentences), or both.^1^ These criteria have since been applied in large cohort studies, including the Centers for Disease Control and Prevention’s Autism and Developmental Disability Monitoring (CDC ADDM) project. Analysis of CDC ADDM data suggested that 26.7% of autistic eight-year-olds would meet criteria for the profound autism designation.^2^

Some have argued that “profound autism” is an “imperative diagnosis,” necessary to ensure that individuals with the highest support needs receive appropriate consideration in both research and public policy.^3^ Others contend that the construct is conceptually flawed and fails to identify a biologically or phenotypically coherent group. The Global Autistic Task Force on Autism Research—a group of autistic professionals and representatives of autistic-led organizations— argued that the term provides little useful information, noting that high support needs arise from many combinations of co-occurring traits and health conditions; they suggest instead using more specific descriptors such as “autistic person with intellectual disability” or “autistic person with minimal language.”^4^ Kapp (2023) similarly noted that, despite acknowledging autism’s “unreliable categorical subtypes,” the Lancet Commission nonetheless proposes a new subtype in profound autism.^5^

A significant concern is that moderate-to-severe intellectual disability and minimal speech do not necessarily co-occur in autistic persons or arise from the same underlying biological mechanisms. From this perspective, profound autism risks becoming a broad catch-all category reflecting severity rather than biology, providing limited explanatory power for research or service provision. Such over-clustering may also hinder policy and clinical decision-making by obscuring meaningful heterogeneity and masking traits that could benefit from more targeted interventions.

Molecular profiling provides a powerful lens through which to test whether the profound autism category represents a distinct biological entity and whether individuals assigned to it constitute a homogeneous group. In the work presented here, we analyze a unique cohort with RNA-seq– derived gene expression and single-nucleotide polymorphism (SNP) data from autistic children and their non-autistic siblings linked to measures of functional impairment. We compare individuals meeting criteria for the proposed “profound autism” construct (PA) with autistic individuals who do not meet those criteria (ASD) to determine whether the PA designation identifies a molecularly distinct population based on gene expression, gene variants, or regulatory architecture (Figure 1).

**Figure 1.**
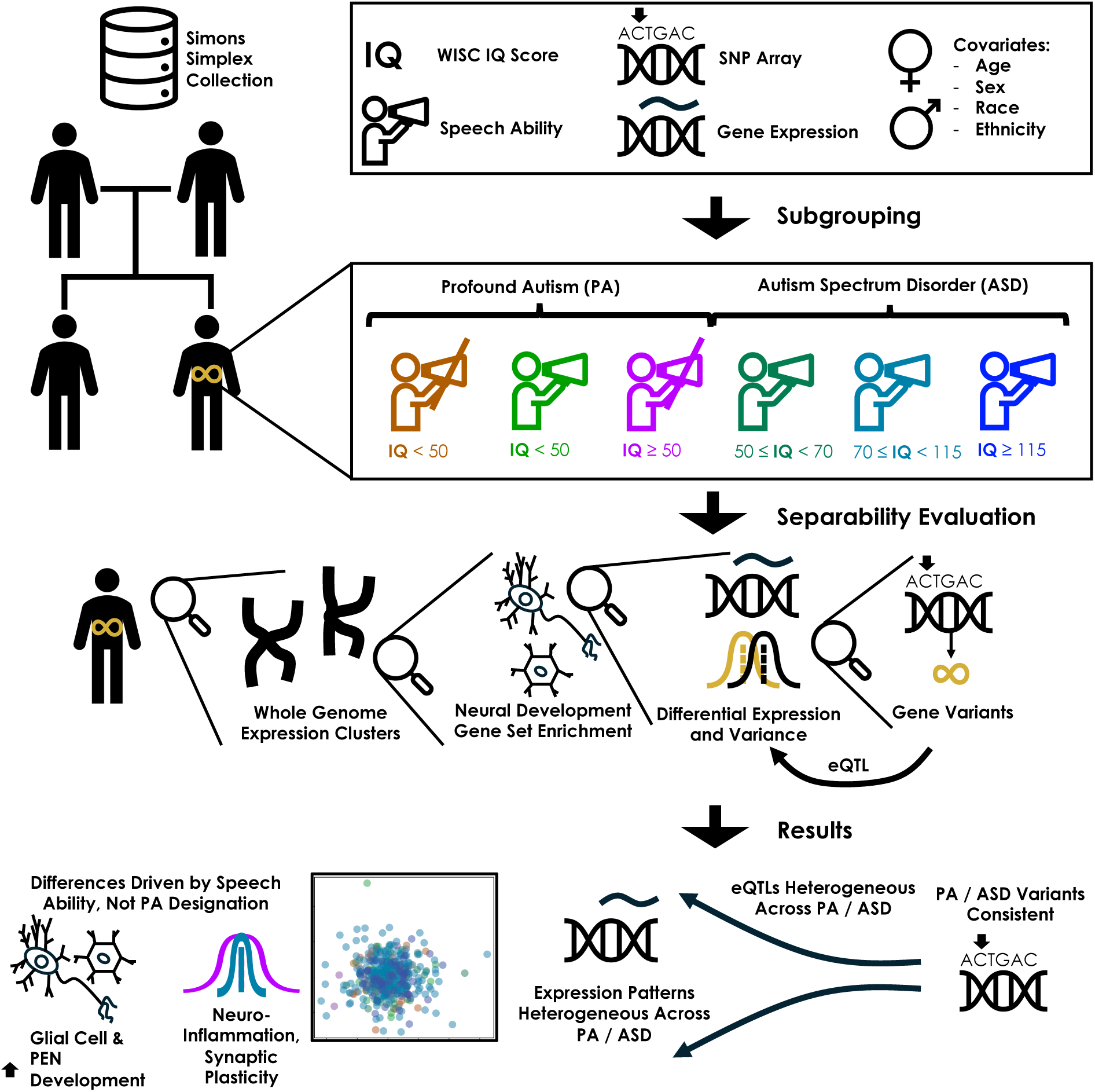
Graphical abstract. Criteria for profound autism were determined using WISC IQ scores and Autism Diagnostic Observation Schedule (ADOS) modules administered to individuals represented in the Simons Simplex Collection (SSC). Participants meeting at least one criterion for profound autism (“Profound Autism / PA”) were separated from autistic children not meeting these criteria (“Autism Spectrum Disorder / ASD”). Individuals in the PA group were further subdivided based on the criteria met while those in the ASD group were divided based on IQ. We compared groups using whole-genome expression profiles, enrichment of neural development gene sets, differential expression and variation analyses, gene variants, and expression quantitative trait loci (eQTLs). Differences were driven primarily by nonspeaking designation rather than PA designation, suggesting that autism-associated SNPs and related factors influence speech-associated pathways differently in nonspeaking versus speaking children.

We tested for gene expression differences between PA and ASD and, across multiple analyses, we found no consistent differences separating these groups. Principal component analysis revealed no systematic separation between groups, even when restricting to proband-specific expression unlikely to reflect shared heritable effects.

We next tested whether specific PA subgroups might differ from ASD in gene sets that had previously been associated with ASD and neuronal processes. Differential expression and variance analyses identified differences only in children meeting the nonspeaking criterion for PA, whereas children meeting the IQ criterion, alone or in combination, showed no consistent differences. Consistent with this finding, enrichment of neurodevelopmental gene sets was driven primarily by speech differences rather than PA designation.

We then assessed whether biological distinctions might instead arise at the level of genetic variants or genetically associated regulatory mechanisms. Autism-associated SNPs were shared across PA and ASD groups, suggesting genetic influences common to both. In contrast, eQTL effects were highly heterogeneous both within PA and between PA and ASD, mirroring the heterogeneity observed in gene expression and failing to define a consistent mechanistic difference.

Together, these results indicate that there is no molecular basis for concluding that PA forms a distinct subgroup of ASD. Instead, the systematic differences we found are primarily associated with speech ability, suggesting that language phenotype may provide a more biologically meaningful axis for stratifying autism.

## Results

To assess the differences between PA and ASD, we used data from the Simons Simplex Collection (SSC), and restricted our analyses to children older than eight years (the age at which the PA designation is generally considered meaningful) who had at least one unaffected sibling; we partitioned individuals into PA and ASD subgroups based on IQ and the Autism Diagnostic Observation Schedule (ADOS) module administered. Throughout this manuscript, we refer to autistic children in both PA and ASD groups as “probands” and their non-autistic siblings as “siblings.” We used IQ < 50 as the cutoff, consistent with DSM-IV criteria for moderate to profound intellectual disability (ID), noting that DSM-V no longer defines ID solely by IQ score.^6^ ADOS modules are selected according to age and speech ability; children assessed with ADOS Module 1 do not demonstrate phrase speech. We therefore used administration of ADOS Module 1 as a proxy for nonspeaking designation. Analyses were restricted to children older than eight years at the time of ADOS administration. To account for developmental differences by age, age was converted into binned variables representing the following ranges in years: [8,10), [10,12), [12,14), [14,16), and [16,18). Additionally, we used Fisher’s exact test to assess the relationship between PA designation and demographic factors, to determine whether demographic biases in diagnosis might confound the analysis. The subgroups and their corresponding significant demographic associations (*p* < 0.05) are presented in Table 1.

**Table 1.**
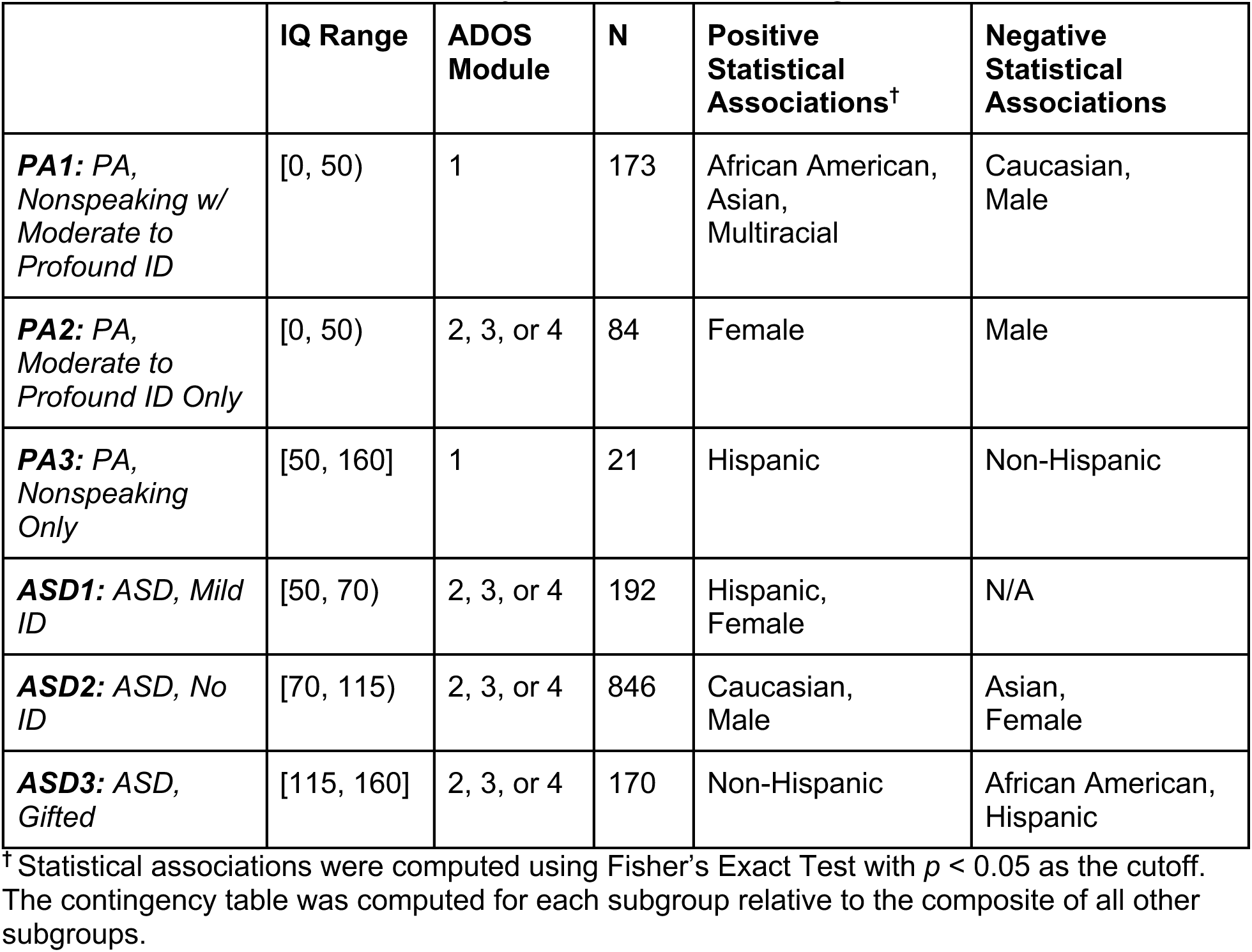
Differences in race, ethnicity, and sex between subgroups.

|  | **IQ Range** | **ADOS Module** | **N** | **Positive Statistical Associations<sup>†</sup>** | **Negative Statistical Associations** |
| --- | --- | --- | --- | --- | --- |
| **PA1:** PA, Nonspeaking w/ Moderate to Profound ID | [0, 50) | 1 | 173 | African American, Asian, Multiracial | Caucasian, Male |
| **PA2:** PA, Moderate to Profound ID Only | [0, 50) | 2, 3, or 4 | 84 | Female | Male |
| **PA3:** PA, Nonspeaking Only | [50, 160] | 1 | 21 | Hispanic | Non-Hispanic |
| **ASD1:** ASD, Mild ID | [50, 70) | 2, 3, or 4 | 192 | Hispanic, Female | N/A |
| **ASD2:** ASD, No ID | [70, 115) | 2, 3, or 4 | 846 | Caucasian, Male | Asian, Female |
| **ASD3:** ASD, Gifted | [115, 160] | 2, 3, or 4 | 170 | Non-Hispanic | African American, Hispanic |

<sup>†</sup> Statistical associations were computed using Fisher's Exact Test with $p < 0.05$ as the cutoff. The contingency table was computed for each subgroup relative to the composite of all other subgroups.

African American, Asian, and multiracial children were overrepresented in the PA subgroup with the most severe impairment (those who are both nonspeaking and have IQ < 50 or *PA1*). Female children were overrepresented in the PA subgroup defined by IQ < 50 alone (*PA2*), while Hispanic children were overrepresented in the nonspeaking PA subgroup with mild or no ID (*PA3*). In contrast, Caucasian and male children were overrepresented in the ASD subgroup without ID (*ASD2*). These patterns are consistent with prior work suggesting barriers to autism diagnosis among racial and ethnic minorities and among women and girls who do not present with severe impairment (Supplementary Tables 1-2).^7–11^ Given these associations, genomic analyses aimed at evaluating profound autism should account for race, ethnicity, and sex to reduce the likelihood of spurious differences in expression or variant analyses.

Having defined PA and ASD subgroups, we evaluated whether there were systematic differences in gene expression, gene variants (SNPs), or regulatory mechanisms (eQTLs) between the subgroups, adjusting for the effects of race, ethnicity, and sex where appropriate. Evaluating the differences between each PA subgroup and ASD allowed us to separate out differences driven by IQ, speech capability, and the combined effects of these. Separating the ASD group into IQ-based subgroups allowed us to further evaluate whether IQ-based differences were specific to those with moderate to profound ID or also persisted across groups with mild ID, no ID, or giftedness.

### Principal Component Analysis Reveals No Systematic Differences Between PA and ASD Along Eigengenes

To assess differences in gene expression between the PA and ASD groups, we first performed principal component analysis (PCA) on gene expression data from all PA and ASD individuals. PCA is a dimensionality reduction technique that transforms a dataset with many potentially correlated variables (such as gene expression levels) into a smaller set of uncorrelated variables called principal components (PC), while retaining as much of the original variation as possible. In our case, each principal component is a linear combination of the original genes based on their expression levels; as such, we sometimes refer to the PCs as “eigengenes” since they represent a weighted sum of genes and their expression levels. In a two-dimensional PC plot, the axes are the first two PCs, which are those that explain the greatest amount of variation in the data. PCA is a widely used technique for exploratory data analysis and has been used in the study of autism to define endotypes or model heterogeneity ^12^. Our goal was to determine whether PA, ASD, and their subgroups represented potential endotypes as evidenced by (1) a clear clustering structure defined by the expression of all of the genes in the genome, or (2) significant differences between subgroups across at least one of the two major principal components (eigengenes), again based on genome-wide expression.

We performed PCA using the *prcomp*() function in R on *logCPM* (log of the counts per million reads from RNA sequencing) genome-wide gene expression profiles, representing the entirety of the 67,073 protein-coding and non-protein-coding genes. To account for expression differences due to sibling sex difference, we performed PCA analysis separately for males and females. As can be seen in Supplementary Figure 1, there is substantial overlap between all PA and ASD groups. As a quantitative assessment, we tested separability between each PA subgroup and the composite ASD group (the combination of Mild ID, No ID, and Gifted ASD subgroups) along each PC using the Wilcoxon rank-sum test with an FDR-adjusted significance threshold of *p* < 0.05 and found no significant separation for either comparison group. This means that gene expression profiles overlapped extensively between PA and ASD groups, indicating that there are no genome-wide transcriptional differences that define a separate PA group. Quantitative results can be found in Supplementary Tables 3-4.

Because the significant overlaps we see between ASD and PA groups could be the result of large numbers of genes that are not directly linked to these subsets but common to everyone, we also tested whether PCA could distinguish find a signal in proband-specific gene expression data. We defined proband-specific gene expression as the standardized difference between affected and unaffected siblings (*logCPM_PSE_ = z*(*logCPM_Proband_*)*– z*(*logCPM_Sibling_*)), evaluating each proband– sibling sex combination: male–male, female–male, male–female, and female–female. As can be seen in Figure 2, there is no significant difference between *logCPM_PSE_* based on this measure, further supporting the conclusion that there is no transcriptional justification for a separate PA subgroup. Quantitative results for can be found in Supplementary Tables 5-8.

**Figure 2.**
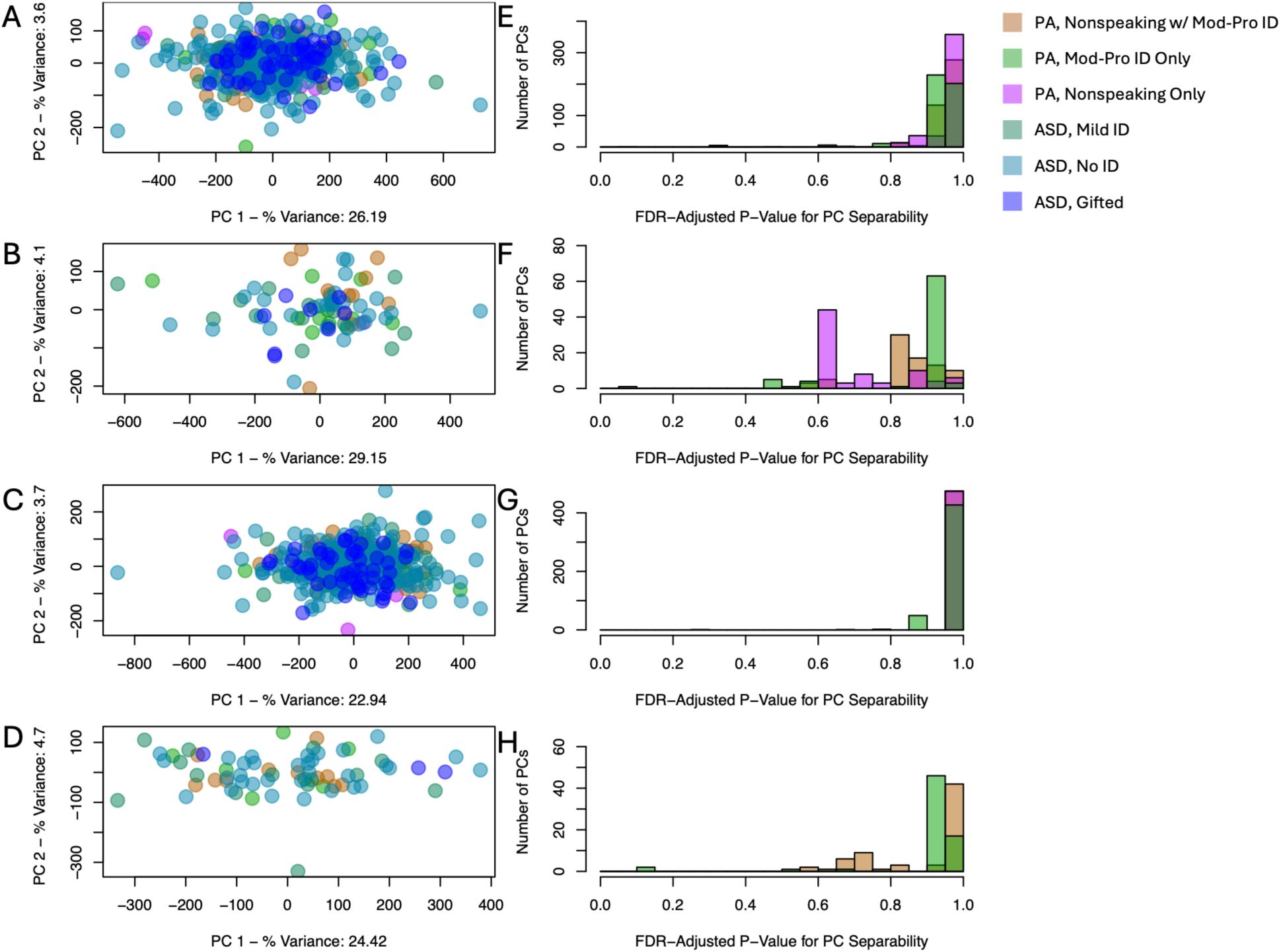
Proband-specific eigengenes do not exhibit systematic differences between PA and ASD. Panels (A-D) illustrate each PA and ASD subgroup projected onto the first two eigenvectors, revealing no meaningful subgroup-specific clustering for(A) male probands with male siblings, (B) female probands with female siblings, (C) male probands with female siblings, and (D) female probands with male siblings. Panels (E-H) illustrate the histogram of FDR-adjusted p-values for a Wilcoxon Rank-Sum Test evaluating separability between each PA subgroup and the combined ASD group along each eigengene, revealing no eigengenes that differ significantly between the PA subgroups and the ASD group in (E) male probands with male siblings, (F) female probands with female siblings, (G) male probands with female siblings, and (H) female probands with male siblings.

### Individual Gene Expression Varies by Speech Ability, Not PA Designation

Although PCA can provide a global picture of potential differences, it may fail to capture the effects of small numbers of genes that might be differentially expressed between subgroups. Consequently, we directly tested for genes that were differentially expressed between PA and ASD subgroups using both *logCPM* gene expression and proband-specific gene expression.

We constructed linear models of each gene’s expression level using Equations 1 for logCPM gene expression levels in probands and 2 for logCPMPSE proband-specific gene expression levels:

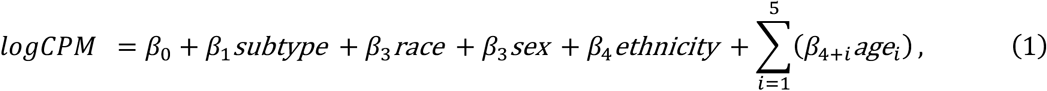

and

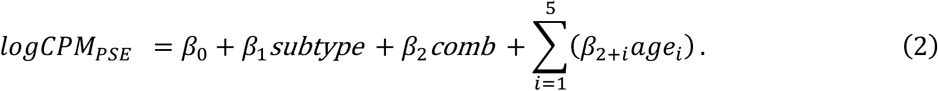

Here, *subtype* represents one of the six PA or ASD subtypes, *comb* represents sex combination (male proband with male sibling, male proband with female sibling, female proband with male sibling, or female proband with female sibling), and *agei* represents age bin *i.* In the *logCPM* evaluations, we included race and ethnicity as covariates. Because all siblings in the data set are full siblings, race and ethnicity covariates were not necessary when evaluating proband-specific gene expression, *logCPM_PSE_*. Significance was determined using *FDR*-adjusted *F*-test *p*-values for *β*_1_.

Consistent with the PCA results, we did not find any genes that were differentially expressed between subgroups using *logCPM* gene expression levels in probands, indicating no consistent PA-specific or moderate to profound ID-specific gene expression. We likewise found no genes differentially expressed between subgroups when evaluating proband-specific gene expression, with the exception of eleven genes that were downregulated in PA3 in comparison to ASD2, two of which were pseudogenes that were also downregulated in PA3 compared to ASD3. In the PA3/ASD2 comparison, two of the genes were protein-coding and have annotated function: *CCL28* and *PXMP2*. Downregulation of *CCL28* has been associated with neuronal loss following prolonged seizures^13^ and with Parkinson’s disease in the striatum, a region implicated in speech motor control.^14,15^. Dysregulation of *PXMP2* has been shown to affect neuronal myelination and contributes to Charcot-Marie-Tooth Disease.^16^ (Supplementary Tables 9-10, Table 2). Although the number of differentially expressed genes is small, the only significant differences we found occur when comparing groups that differ in speech ability, not in PA designation.

**Table 2.** Summary of the number of genes found to be differentially expressed (*logCPM_PSE_*) in comparing PA and ASD subgroups.

|  | PA1 | PA2 | PA3 | ASD1 | ASD2 | ASD3 |
| --- | --- | --- | --- | --- | --- | --- |
| PA1 | -- | 0 | 0 | 0 | 0 | 0 |
| PA2 | -- | -- | 0 | 0 | 0 | 0 |
| PA3 | -- | -- | -- | 0 | **11** | **2** |
| ASD1 | -- | -- | -- | -- | 0 | 0 |
| ASD2 | -- | -- | -- | -- | -- | 0 |

Although differences in the level of gene expression between states are the most widely used measure of molecular differences between phenotypic states, differences in the variation of genes within populations can also provide insight into the functional drivers of biological processes ^17^. The rationale is that genes that play important roles in defining biological states are more consistently regulated across individuals and that degradation of regulatory control can result in phenotypic change. We tested whether any of the PA subgroups exhibited altered patterns of variance in comparison to ASD (and therefore represent a distinct transcriptomic subgroup within the broader construct of ASD) using Levene’s test. We compared gene expression variance in each PA subgroup to gene expression variance across the composite ASD group (the combination of Mild ID, No ID, and Gifted ASD subgroups), considering both the *logCPM* and *logCPM_PSE_* values defined in the previous section. We compared the PA subgroups to the composite ASD group here rather than the ASD subgroups because the goal was to determine whether gene variance within each PA subgroup differed from broader ASD, not from subsets of the ASD population. Statistical significance was defined as FDR-adjusted *p* < 0.05.

We did not find significant differences in variance between ASD and either PA1 or PA2 (those subgroups with intellectual disabilities, independent of speech impairment). However, when comparing ASD and PA3, we found significant increases in the variance of 103 genes and five genes with decreased variance. The result was even more striking when comparing proband-specific expression where we found 356 genes with increased expression variance in PA3 and none with decreased variance (Table 3, Supplementary Tables 11-12).

**Table 3.** Significant Gene Counts for logCPMPSE and logCPM Variance Analysis.

|  |  | *Increased Variance Gene Count* | *Decreased Variance Gene Count* |
| --- | --- | --- | --- |
| $\log CPM$ | PA1 | 0 | 0 |
|  | PA2 | 0 | 0 |
|  | PA3 | **103** | **5** |
| $\log CPM_{PSE}$ | PA1 | 0 | 0 |
|  | PA2 | 0 | 0 |
|  | PA3 | **356** | 0 |

As a way of exploring the functional consequences of this increased variance, we performed gene set enrichment analysis (GSEA). We used the *fgsea* R package (version 1.32.4)^18^ with pathways from MSigDB ()^19^ and defined the input scores as described in Equation 3, where *p*_j_ is the *p*-value for gene *j* returned from Levene’s Test, *σ*^2^*_PA_*_3,j_ is the variance of *j* in PA3, and *σ*^2^*_ASD,j_* is the variance of *j* in ASD.

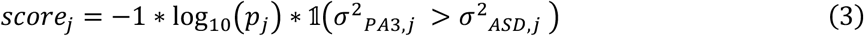

The *fgsea* method uses genes ranked based on some statistical measure and then tests for gene sets that are significantly overrepresented at the extreme of the distribution. Gene sets enriched among the 103 significant genes that exhibited increased *logCPM* variance in PA3 compared to ASD included those in pathways related to myelination (phosphatidylinositol pathways),^20^ neuroinflammation (*IRF3*^21^ and *NF-κB* pathways^22^ and antigen processing),^23^ and synaptic plasticity (integrins^24^ and metabotropic glutamate receptors).^25^ These results suggest greater variability in these biological processes among nonspeaking autistic children with mild to no ID compared with speaking autistic children with mild to no ID (Supplementary Tables 13-14). Although altered neuroinflammation, myelination, and synaptic plasticity have been reported in autism compared with non-autistic controls,^26–28^ the variability of gene expression within these pathways has not been examined in nonspeaking autistic children specifically, to our knowledge. Some of these pathways overlap with other neurological conditions. *IRF3* has been linked to Alzheimer’s disease (AD),^21^ a condition for which autistic individuals and their families may show increased risk,^29^ and 17q12 copy-number variation (CNV) syndrome, associated with autism and speech delay.^30^

When analyzing the 356 genes showing significant increases in variance in proband-specific gene expression, we found enriched antigen processing, *NF-κB*, *IRF3*, and 17q12 copy-number variation syndrome pathways, similarly to the *logCPM* variance analysis. These results indicate that not only do neuroinflammatory processes and AD and 17q12-related biological processes exhibit greater variability in PA3 compared to ASD, but that they do so with respect to genes expressed differently from the siblings of children with PA3, suggesting that individual-specific factors contribute to differential regulation of these pathways in addition to regulatory processes shared between siblings. Pathways related to other neurological conditions include Aβ-related alterations in speech prosody within AD^31^ and 8p23 CNV syndrome, also associated with speech delay.^32^ Increased variance of genes related to Aβ alterations and 8p23 CNV syndrome in PA3 *logCPM_PSE_* suggest individual-specific regulation of these mechanisms.

Collectively, these results indicate that the PA3 non-verbal group’s increased gene expression variability relative to ASD in processes related to neuroinflammation, myelination, and synaptic plasticity. Of these, neuroinflammatory processes appear to exhibit individual-specific regulation leading to increased variability, whereas the variability in myelination and synaptic plasticity pathway activity is related to gene regulatory processes shared between siblings. Further, increased variability exists in genes related to other neurological conditions linked to speech challenges, including AD, 17q12 CNV syndrome, and 8p23 CNV syndrome, of which AD and 8p23 syndrome exhibit some degree of individual-specific regulation. In combination with the consistent downregulation of *CCL28* and *PXMP2* and their respective roles in neuronal loss, speech motor control, and myelination, the high variability of pathways associated with other neurological conditions and other myelination, neuroinflammation, and synaptic plasticity pathways in PA3 indicate that the most meaningful distinction between subgoups of individuals with ASD may be associated with language.

### Speech Ability, Not PA Designation, Drives Enrichment of Heterogeneity-Related Gene Sets

Given that the only meaningful differences in gene expression levels and variance occurred when comparing individuals with PA3 to those with ASD, we tested whether gene sets previously linked to ASD were also significantly associated with differences in speech, IQ, or a combination thereof. We assembled gene sets based on a meta-analysis performed by Litman et al. that linked *de novo* loss-of-function variant gene sets to phenotypically-defined subgroups within autism^33^; we downloaded these from their GitHub repository (https://github.com/FunctionLab/asd-pheno-classes/tree/main/GenomicAnalyses/gene_sets). The sets from Litman that we chose included the following:

- Autism-associated genes (Sanders et al.)^34^
- Genes associated with autism or developmental delay (Satterstrom et al.)^35^
- Fragile X Mental Retardation Protein (FMRP) targets^36^
- Genes with high evolutionary constraint
- Genes associated with the postsynaptic density,^37^ a region that regulates synaptic plasticity, that were obtained from the Genes to Cognition (G2Cdb) database^38^
- Autism-associated genes from SFARI Gene 2.0^39,40^
- Genes associated with specific neurodevelopmental trajectories in principle excitatory neurons, inhibitory interneurons, and glial cells,^41^ – trajectories included an increase from fetal to adult stages (*Up*), a decrease from fetal to adult stages (*Down*), an increase in fetal and adult stages only (*Osc Up*), and an increase from infancy to adolescence only (*Osc Down*)

To evaluate enrichment of each of these gene sets based on differences in gene expression, we used the *fgsea* R package, defining each gene set as an ASD-associated pathway. The genes were ranked based on the score defined in Equation 4.

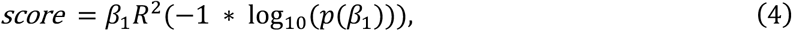

where *R*^2^ is determined by the fit of the linear model used for gene expression analysis, defined in Equation 2. This allowed us to account for *p*-value, effect size, and goodness-of-fit as factors in determining gene score. We limited our analysis to the proband-specific gene expression because Litman *et al*., focused on proband-specific *de novo* gene variants, which we expect to affect proband-specific gene expression independently of sibling gene expression.

We performed comparisons to evaluate the roles of speech ability only, IQ only, and combined PA designation (represented by PA1) as follows:

1. **IQ(1-4):** Comparisons between children in differing IQ groups, including PA1 and PA3 (**1**) PA2 and the ASD1 (**2**), ASD1 and ASD2 (**3**), and ASD2 and ASD3 (**4**).
2. **Speech(1-4):** Comparisons between nonspeaking children and speaking children, including between PA1 and PA2 (**1**), PA3 and ASD1(**2**), PA3 and ASD2 (**3**), and PA3 and ASD3 (**4**).
3. **Comb(1-3):** Comparisons between PA1 and the ASD groups, i.e., (**1**) ASD1, (**2**) ASD2, and (**3**) ASD3.

Because ties in gene score rank can affect fgsea outcomes, we ran each analysis ten times with a random seed and reported the average signed log-scaled FDR-adjusted p-values, where the sign corresponded to the direction of enrichment in GSEA.

Most gene sets showed higher expression in **Speech** comparisons than in **Comb** comparisons, particularly between PA3 and ASD3 (**Speech4**). The persistence of these differences even within PA-designated individuals in the comparison between P1 and P2 (**Speech1**) indicates that nonspeaking designation has a stronger association with gene set upregulation than PA designation, defined broadly as the composite PA group or narrowly as P1 only. The **IQ1** comparison (PA1 and PA3) likewise showed downregulation for many gene sets (significance cutoff is 1.30, a negative log-scaled FDR-adjusted *p*-value of 0.05), indicating differences in gene set enrichment between persons who are nonspeaking due to factors other than ID and those who may be nonspeaking in part due to ID (Supplementary Figure 2, Supplementary Table 15).

To limit the influence of gene set overlap on enrichment results, we repeated the *fgsea* analysis with each gene set limited to the set of genes unique to that set (Supplementary Table 16).. We found that several gene sets were similarly enriched in the Speech comparisons, especially **Speech1** and **Speech4**, again indicating a stronger association between nonspeaking designation than between broadly or narrowly defined PA designation. The significance of enrichment was highest for SFARI autism genes, developmental delay genes from Werling et al., glial cell genes with increasing expression in fetal and adult stages, and principal excitatory neurons with increasing expression from fetal to adult stages (Figure 3, Supplementary Table 17).

**Figure 3.**
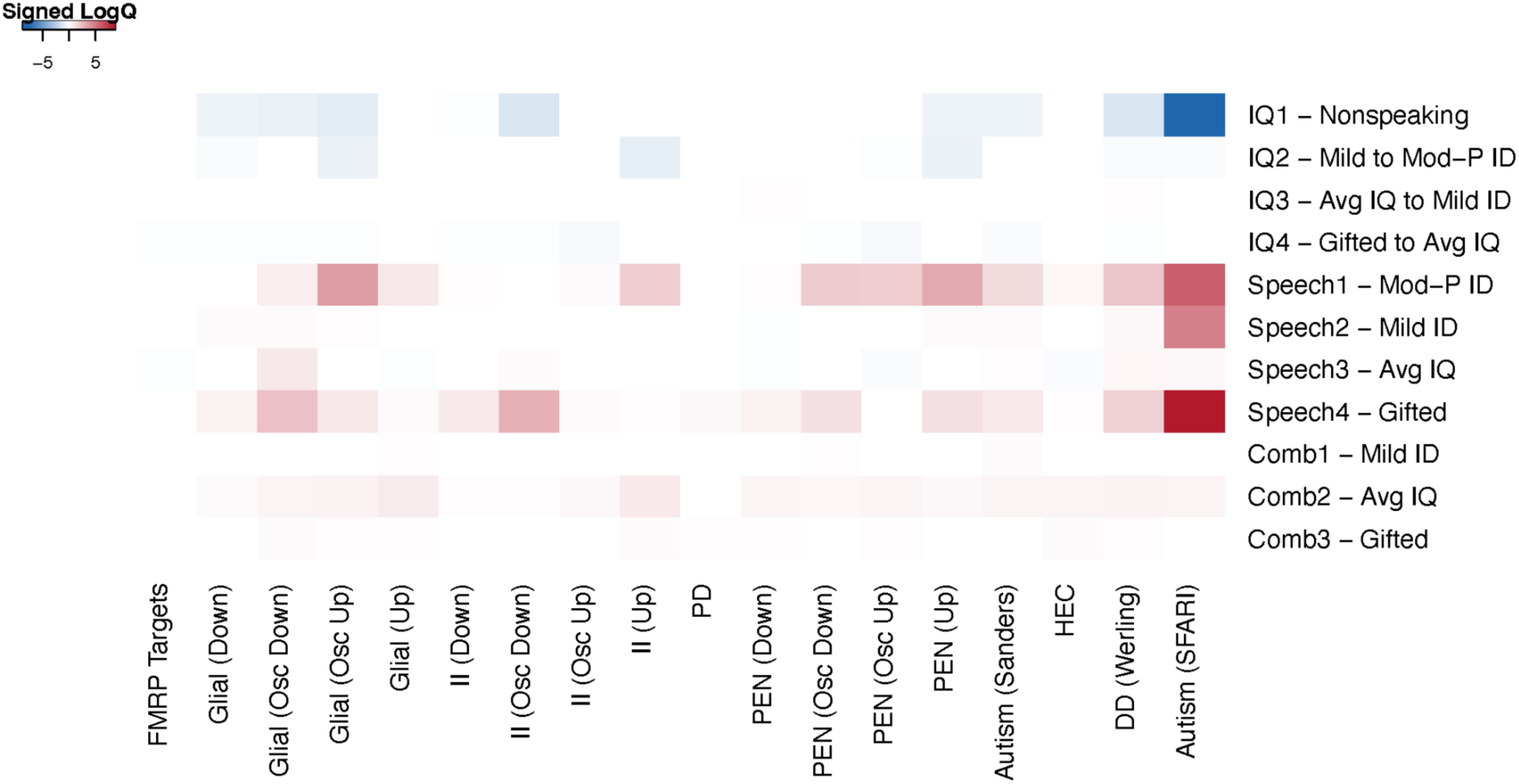
Enrichment Analysis Heatmap for Genes Associated with Autism, Developmental Delay, or Neuronal and Glial Cell Development. Rows labeled IQ represent comparisons between groups with the same speaking designation but different IQ ranges. Rows labeled **Speech** represent comparisons between speaking and nonspeaking children within the same IQ range. Rows labeled **Comb** represent comparisons between groups that differ in both IQ range and speaking designation. Colors represent the signed log10(q-value) after FDR adjustment, with negative enrichment shown as negative values. Mod-P ID refers to moderate-to-profound intellectual disability.

These various analyses that compare features of gene expression between PA and ASD subgroups lead to one conclusion: there are no convincing expression differences that support the designation of profound autism as a molecularly distinct subgroup. At best, there may be some difference between individuals with ASD and those assigned to the PA3 (that is, between nonspeaking and speaking children with an IQ ≥ 50); we see similar differences between the PA1 and PA2 subgroups (children with IQ < 50, differentiated based on speech), which suggests that transcriptomic differences may drive speech ability independently of intellectual disability.

### Autism-Associated SNPs Are Consistent Between PA and ASD

Although we tend to think of gene expression differences as being close to phenotypic differences, the expression profiles we collect are typically representative of a static point in time and so could miss factors that drive different developmental trajectories that could distinguish PA and ASD or alter function in other ways. Many gene variants have been associated with autism^42^ and so we tested whether these might be differentially represented in the PA and ASD subgroups.

We performed a genome-wide association study (GWAS) using date from the SSC to test whether autism-associated genetic variants (single nucleotide polymorphisms; SNPs) were more common in the PA or ASD relative to the general population than would be expected by chance. We tested all 1,072,814 nuclear SNPs available from SSC for overrepresentation in each of the following subgroups of autistic children: (1) the ASD composite subgroup, comprised of ASD1, ASD2, and ASD3, (2) the mild-to-no-ID ASD subgroup, comprised of ASD1 and ASD2, (3) the no-ID ASD subgroup, comprised of ASD2 and ASD3, (4) the PA composite subgroup, comprised of PA1, PA2, and PA3, (5) the PA nonspeaking subgroup, comprised of PA1 and PA3, and (6) the PA moderate-profound ID subgroup, comprised of PA1 and PA2. Analyses 2, 3, 5, and 6 were performed to ensure that analyses 1 and 4 were not heavily influenced by the smaller subgroups of gifted children, speaking children with mild ID, or children meeting only one criterion for PA. We took this leave-subgroup-out approach rather than evaluating each PA and ASD subgroup separately because of the large sample size requirement for GWAS and the small sample sizes of some of the subgroups.^43^ For each GWAS, we compared the subgroups of probands to their non-autistic siblings and parents, with autism diagnostic status (autistic/non-autistic) as the outcome. Covariates included the top six principal components (PCs) from SNP data as *ancestry PCs* (Supplementary Figure 3).

We considered those SNPs associated with autism (FDR-adjusted *p* < 0.05) to be nominally significant. This is far below the threshold normally considered significant in GWAS studies, but it allowed us to at least identify possible associations and rule out many more. Details of the GWAS analysis are provided in the Methods. Using these criteria, we identified 92 nominally significant SNPs in the combined ASD group and 76 in the combined PA group; the 76 PA SNPs were a proper subset of the ASD SNPs (Figure 4, Supplementary Table 18).

**Figure 4.**
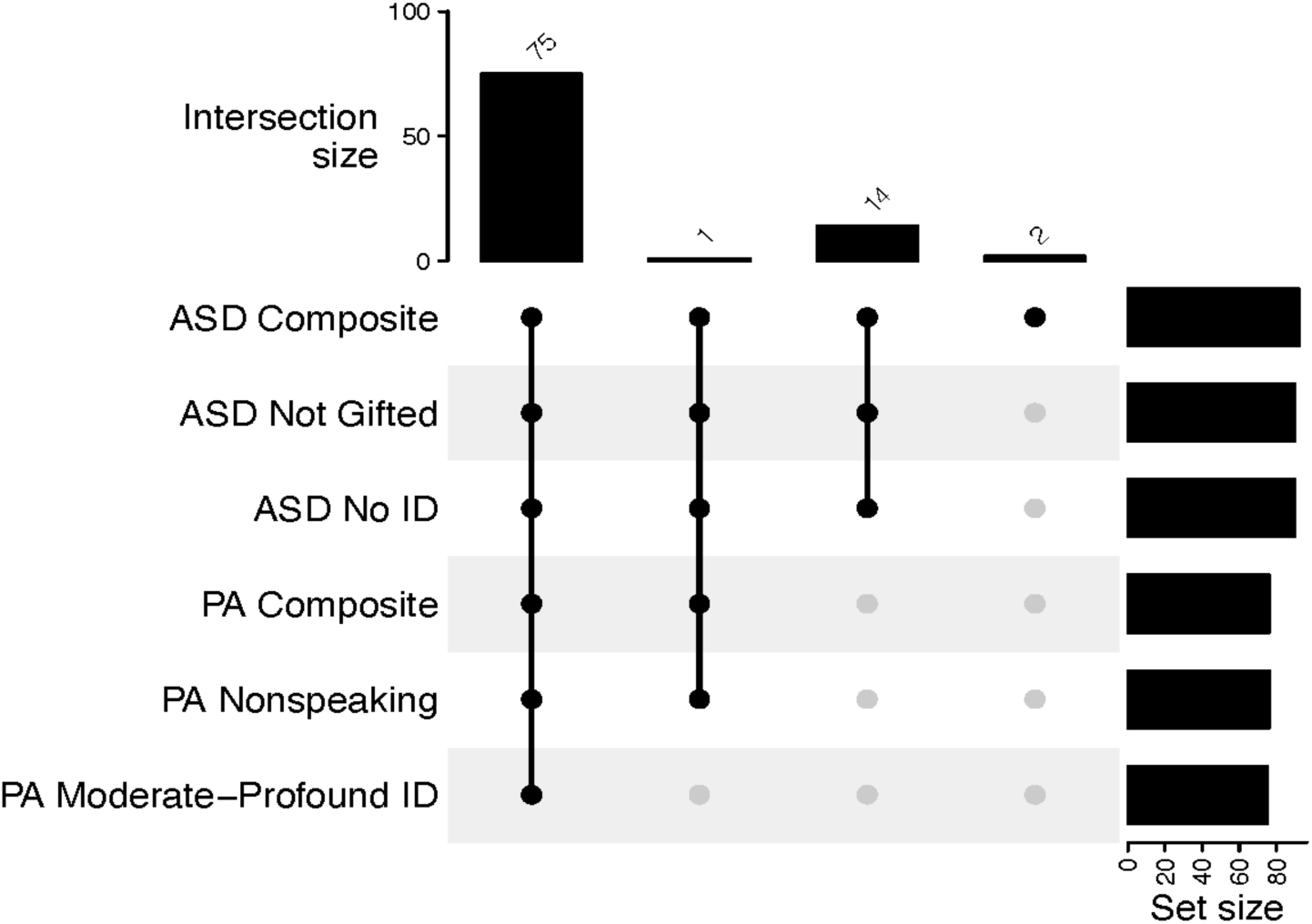
Nominally significant SNPs Overlapping Across Group-Specific GWAS. In this UpSet plot, “Set size” refers to the number of significant SNPs from each group, and “Intersection size” to the number of significant SNPs in each intersection. A solid black circle indicates that a group is part of an intersection, and a solid gray circle indicates that it is not.

For the 16 SNPs that were found to be nominally significant only in ASD but not in PA, we mapped them to genes using coordinates from GENCODE Human Release 49, which we converted to BED format using BEDOPS (v2.4.41) *gtf2bed*.^44^ Each SNP was assigned to the nearest gene as a way of understanding their potential functional roles (Table 4). Several of the genes linked to the ASD-only nominally significant SNPs have been linked to autism. *SHANK2* regulates synapse development and has been linked to autism in *in vitro*, *in vivo*, and clinical studies.^45^ A protein-protein interaction module including *RGS7*, a regulator of GTPase activity, was found to be enriched in autistic children in comparison to non-autistic children by Topal et al.^46^ *PCDH11X*, which is involved in nervous system development, has been associated with autism in comparison to non-autistic controls in multiple cohorts.^47^ *ADARB1*, which encodes an enzyme that edits the glutamate receptor within the nervous system, has been proposed as a biomarker for ASD,^48^ and possible differences in the metabolism of riboflavin (facilitated by *RFK*) have been linked to ASD.^49^ These results illustrate that a variety of genes involved in neurological function and development, as well as metabolism, are associated with autism whether or not the criteria for PA are met. Further, 80 of the 92 SNPs we found in this GWAS analysis are from the pseudo-autosomal region of the X chromosome (PAR), consistent with the hypothesis of X chromosome involvement in autism^47^ – again, our results illustrate that this involvement is likely consistent across those meeting and not meeting the criteria for PA. Importantly, we did not find any SNPs that were associated specifically with PA and not with ASD.

**Table 4.** PA p-value rankings of SNPs nominally significant only in ASD and their associated genes.

| Chromosome | Gene Name | SNPs<sup>‡</sup> | Scaled Ranking in Profound Autism (0-1) |
| --- | --- | --- | --- |
| 1 | RGS7 | SNP1-241147595,<br>rs7533316 | 3.16e-04,<br>7.08e-03 |
| 3 | RFTN1 | SNP3-16382477,<br>SNP3-16382523,<br>SNP3-16383255,<br>SNP3-16383727,<br>SNP3-16454562 | 3.20e-05<br>3.24e-05<br>3.28e-05<br>3.16e-05<br>3.32e-05 |
| 9 | Pseudogene Similar to Part of a Novel Protein KIAA1688 | SNP9-31110120 | 4.63e-05 |
| 11 | SHANK2 | SNP11-70539974 | 3.40e-05 |
| 12 | ATP2A2 | SNP12-110351451 | 3.52e-05 |
| X (PAR) | ADARB1 Pseudogene | rs2984684 | 3.44e-05 |
|  | RFK Pseudogene | rs5941380 | 1.93e-03 |
|  | SERBP1P4 | rs35970878,<br>rs35429716 | 1.18e-04,<br>3.85e-05 |
|  | CTAG1A | rs1084422 | 3.77e-05 |
|  | COPZ1 Pseudogene | rs306934 | 5.66e-05 |

‡ The underlined SNPs were nominally significant only in ASD rather than PA. “PAR” refers to the pseudo-autosomal region of sequence similarity between chromosomes X and Y.

Although these 16 SNPs were found to be nominally significant only in ASD and not in PA, we examined their ranking in the PA analysis. Although they did not reach our threshold for nominal significance, they fell within the lowest 1% of *p*-values in the PA group (meaning that they were more strongly associated with PA than 99% of SNPs evaluated). This indicates that they had a strong association despite not reaching significance with PA despite not meeting our threshold. This is not surprising, however, given the much larger number of individuals in the ASD group compared to the PA group (1,208 vs 278). This, once again, demonstrates a lack of molecular support for the PA designation.

### Regulatory Heterogeneity Persists Across PA Subgroups and Random PA Subsets

SNPs can also exert regulatory effects by altering the binding of transcription factors that activate or repress gene expression^50,51^. We performed expression quantitative trait locus (eQTL) analyses in all PA and ASD subgroups to test for associations between genetic variants and the expression levels of each gene. We performed eQTL analysis on probands for whom both SNP and gene expression data were available, including ancestry PCs, sex, and binned age as covariates. We tested for both cis- and trans-eQTLs within each PA and ASD subgroup; additional details on the eQTL analysis are provided in the Methods. The significant differences in the number of eQTLs found in the ASD and PA analyses again represent sample size effects.

We compared the sets of eQTLs calculated for both the proband *logCPM* and on proband-specific gene expression, *logCPM_PSE_* and found minimal overlap between those identified in the PA and ASD subgroups – a result that would also be expected for randomly selected subgroups in a heterogeneous population due in part to biological heterogeneity. Biological heterogeneity is expected in random subsets because eQTLs are known to vary widely across individuals and contexts, explained in part by differences in transcription factor binding affinity and chromatin accessibility, as well as additional genetic factors that vary between individuals^52,53^. This result suggests heterogeneity in eQTLs between children meeting and not meeting the criteria for PA as well as between subgroups of PA and subgroups of ASD. eQTL consistency between PA subgroups appears no greater than eQTL consistency between PA and ASD—the number of eQTLs shared between PA1 and PA2 was lower than the number of eQTLs shared between PA1 and ASD2. This result was echoed in both the *logCPM* and proband-specific *logCPM_PSE_* analyses (Figure 5).

**Figure 5.**
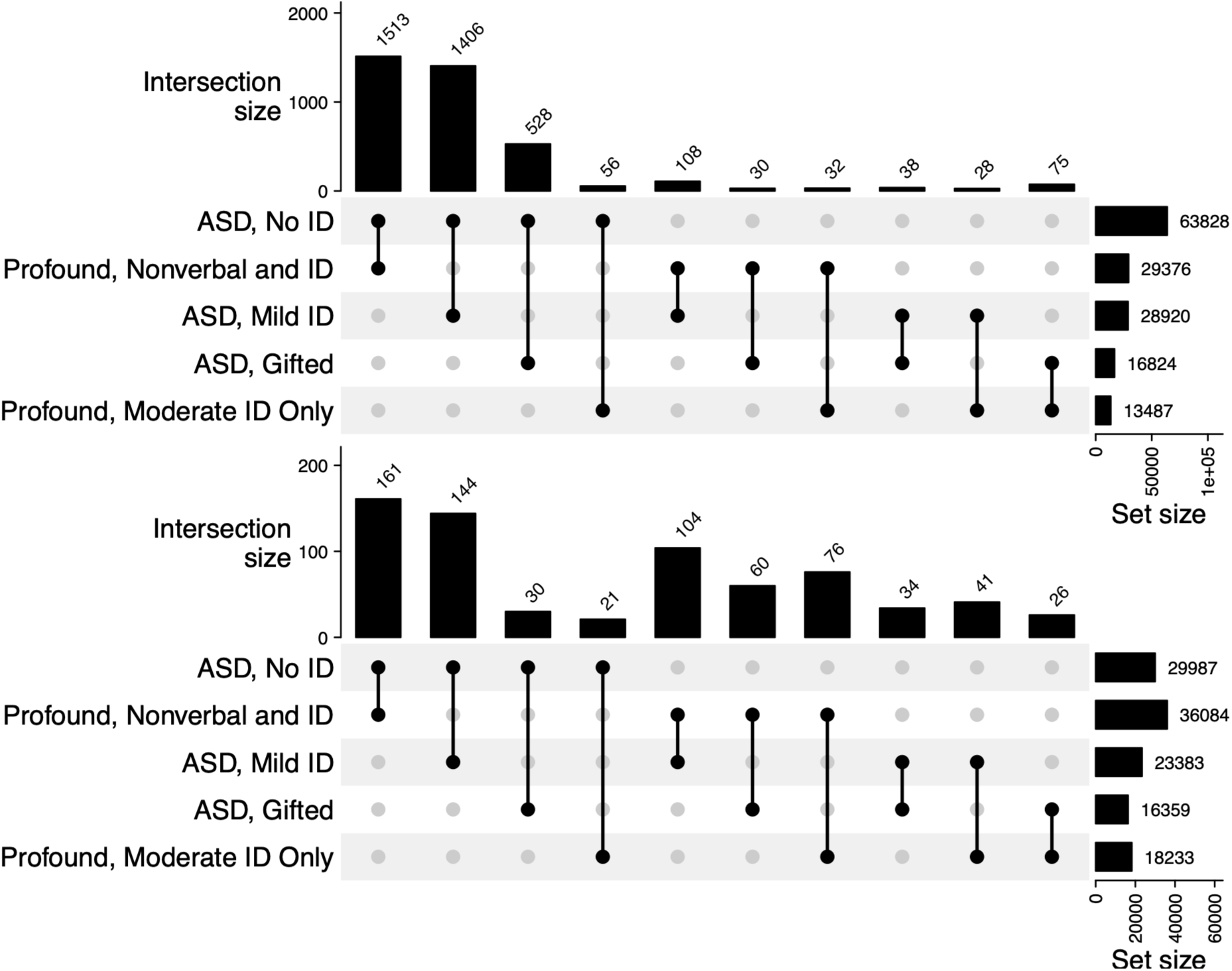
Significant cis-eQTLs Overlapping Across Subgroup-Specific Analyses. These results include both eQTLs computed with *logCPM* gene expression (top) and proband-specific gene expression, *logCPM_PSE_* (bottom). In this UpSet plot, “Set size” refers to the number of eQTLs from each group, and “Intersection size” to the number of significant eQTLs in each intersection. A solid black circle indicates that a group is part of an intersection, and a solid gray circle indicates that it is not. In this plot, the number of eQTLs for each intersection of two PA/ASD groups includes those found in additional groups.

To determine whether the inter-group heterogeneity we observed reflected true biological differences between groups or heterogeneity among individuals with respect to PA, we investigated to what extent eQTLs overlapped between random subsets of children in PA1. To conduct this analysis, we randomly split samples within PA1 into two subsets, then independently inferred cis-eQTLs and trans-eQTLs within each subset, across ten independent runs. We observed minimal overlap in eQTLs between subsets, illustrating that not only do eQTLs not persist between PA1 and PA2, but that eQTLs are also highly variable within PA1, echoing the high heterogeneity within this group observed in our gene expression analyses (Table 5). The Jaccard overlap is described as the percentage of eQTLs overlapping the two random sets, described in the Methods.

**Table 5.**
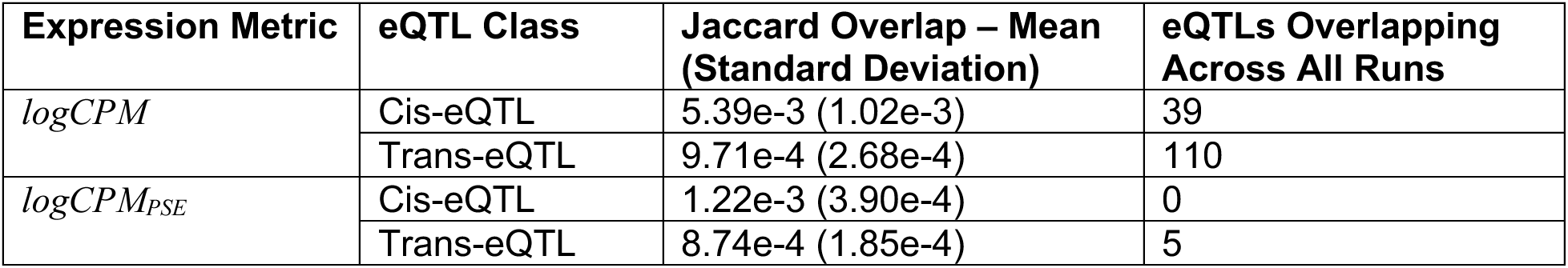
eQTL overlap between randomized subsets of PA1.

| Expression Metric | eQTL Class | Jaccard Overlap – Mean (Standard Deviation) | eQTLs Overlapping Across All Runs |
| --- | --- | --- | --- |
| *logCPM* | Cis-eQTL | 5.39e-3 (1.02e-3) | 39 |
|  | Trans-eQTL | 9.71e-4 (2.68e-4) | 110 |
| *logCPM<sub>PSE</sub>* | Cis-eQTL | 1.22e-3 (3.90e-4) | 0 |
|  | Trans-eQTL | 8.74e-4 (1.85e-4) | 5 |

While the low eQTL overlap between random subsets is indicative of inter-group heterogeneity, it is also possible that many eQTLs are spurious due to small sample sizes (60 PA1 samples included both SNP and gene expression data). To determine the PA specificity of higher confidence eQTLs, we examined the subset of eQTLs overlapping across all independent runs, finding that none were PA-specific in the subgroup-based eQTL analyses and suggesting that there is no regulatory mechanism exclusive to PA. These pairs are further described in Supplementary Table 19.

The heterogeneity of eQTLs among autistic children suggests that variability in gene regulation contributes to broad expression heterogeneity despite shared autism-associated SNPs across groups. Moreover, heterogeneity persists across PA subgroups, PA and ASD subgroups, and randomized subsets of PA1, illustrating that even under a narrow definition of PA, the regulatory landscape is highly heterogeneous.

## Discussion

Subgroup classification remains one of the most contentious issues in autism research and service provision. In 2013, the American Psychiatric Association (APA) consolidated three previously distinct autism spectrum diagnoses in the Diagnostic and Statistical Manual of Mental Disorders, Fourth Edition (DSM-IV) into a single unified Autism Spectrum Disorder category in DSM-5. In advocating for a unified diagnosis, the APA and other supporters noted that prior literature documented inconsistent and often arbitrary application of the earlier diagnostic categories, and that separating individuals into multiple diagnoses often complicated rather than clarified research and service provision.^54,55^ In a memorable phrase, the APA described the DSM-IV criteria as “equivalent to trying to ‘cleave the meatloaf at the joints.’” Instead, the organization argued that autism “is best represented as a single diagnostic category that is adapted to the individual’s clinical presentation by inclusion of clinical specifiers.”^56^

This was not without controversy, raising concerns that the ‘lumping’ of different groups of autistic persons into a single diagnosis might fail to acknowledge “distinct biological substrates” and give insufficient attention to persons with the most severe impairment.3,55 The proposed “profound autism” construct proposed in 2022 and subsequent attempts to establish it as a diagnostic category constitutes an attempt to turn back the clock and reintroduce subclassifications into autism.

It is important to distinguish the broader question of whether subclassifications may be useful from the specific proposal of “profound autism.” Any classification must be evaluated according to the scientific, clinical, and social utility it provides in a particular context. Our findings indicate that, in the context of genetic and genomic research, “profound autism” does not represent a distinct biological substrate and does not offer explanatory value for this type of investigation. This conclusion does not rule out the possibility that the construct might provide value in other settings, such as service provision or other forms of research. However, proponents have not yet presented evidence demonstrating such utility.

Researchers, advocates, and policymakers considering adoption of the “profound autism” construct should therefore proceed with caution. Classification systems can produce unintended harms. Just as excessive “lumping” can obscure meaningful differences, excessive “splitting” can create the false impression that the diversity of autism has been subdivided in a biologically meaningful way when the evidence does not support such precision. Establishing a separate diagnostic category may also imply that advances relevant to one population do not apply to another.

This concern is particularly acute given how closely the battle lines in debates surrounding “profound autism” closely mirror longstanding conflicts among stakeholders regarding how best to allocate autism research dollars, the appropriateness of institutionalization and similar congregate residential service provision models, the legality of sub-minimum wage employment, and other hot-button conflicts in autism advocacy.57,58 Insofar as “profound autism” seeks to establish a category for the purpose of garnering advantage in these debates, it functions more as a political rather than a scientific classification.

Recently, other, more nuanced subclassifications for better understanding autism based on behavioral phenotypes have been proposed, supported by evidence that they correspond more closely to underlying genetic profiles.^33^ Similarly, our findings highlight genetic commonalities among autistic individuals who do not speak but have mild or no intellectual disability, suggesting that it may be more informative to characterize heterogeneity in autism by reference to specific functional traits rather than attempting to subdivide the multidimensional construct of autism itself. This approach not only aligns with the DSM-5 working group’s recommendation to use clinical specifiers and with recent recommendations from the Global Autistic Task Force, but also with our understanding of biological processes. Although there may be a range of traits, such as speech impairment or intellectual disability, the functional mechanisms involve distinct neurological processes and the activation of distinct molecular pathways in brain cells.

At the same time, an important gap in our understanding remains. We do not yet know the degree to which autistic individuals with specific impairments resemble individuals with other developmental disabilities who share the same characteristics. The functional impairments included in the “profound autism” construct occur outside autism as well. A substantial body of research has documented diagnostic substitution, in which individuals who might previously have received a diagnosis of intellectual disability are now diagnosed with autism instead of, or in addition to, ID.^59,60^ Future work should therefore examine whether severe speech or cognitive impairment within autism represents a distinct phenomenon or reflects mechanisms shared with other developmental conditions. Although prioritizing research and services for individuals with severe impairments is both important and overdue, it remains unclear whether an autism-specific framework is the most informative way to pursue that goal.

The creation of a new diagnostic category, profound autism, for individuals with autism spectrum disorder, should be grounded in meaningful biological differences rather than arbitrary classifications. Using the diagnostic criteria for profound autism, we searched one of the largest available comprehensive molecular profiling datasets, the Simons Simplex Collection, for robust features that would distinguish it from other sub-categories of the disorder. We tested various measures of gene expression, representation of genetic variants, and the intersections between genetic variants and the genes they appear to regulate. Despite this extensive search, we found no robust molecular basis for a separation between individuals separated by a PA or ASD classification, whether taken as a cohesive group or when considering more granular divisions based on intellectual disability or speaking ability.

As a whole, our analyses suggest that, if anything, it is more meaningful to study differences between speaking and nonspeaking autistic persons, particularly when considering interventional strategies. Speech-related gene expression differences are most prominent in the absence of moderate-to-profound intellectual disability, suggesting that nonspeaking individuals whose speech challenges are not driven by intellectual disability may require more targeted intervention strategies focused on speech mechanisms. Indeed, it may be useful to prioritize research into interventional strategies for individuals falling into this narrowly defined class.

We note some limitations to our analyses. First, our analysis is limited to children between the ages of eight and eighteen years old and therefore excludes data from adults—to our knowledge, no data set currently exists for autistic adults that includes gene expression data, gene variant data, and phenotypic data sufficient to characterize subjects as meeting the criteria for profound autism. Further, particularly for the GWAS and eQTL analyses, the sample size is smaller than necessary to find robust positive signals and the sample size imbalance presents challenges in doing direct comparisons between stratified analyses based on PA/ASD classifications. Concerns about sample size and statistical power prevented us from including sex or independently analyzing the less prevalent ASD and PA subgroups in our GWAS analyses. Despite these limitations, our results present a compelling arguments against the creation of one or more PA subgroups. This results in direct consequences not only for the clinical definitions presented in the Diagnostic and Statistical Manual of Mental Disorders (DSM), but also for public policy decisions that rely on understanding subgroup classifications.

## Methods

### Simons Simplex Collection

The Simons Simplex Collection (SSC) includes data from 2,643 families in which exactly one child has been diagnosed with autism (referred to as the proband). These data include phenotypic measures for probands, mRNA-seq from lymphoblastoid cell lines (LCLs) for probands and siblings, and single-nucleotide polymorphism (SNP) data from LCLs for probands, siblings, and parents. Phenotypic measures include demographic information, medical history, Autism Diagnostic Observation Schedule (ADOS) item-level and composite scores, and data collected from other diagnostic instruments.^61^

We requested access to this resource through the Simons Foundation Autism Research Initiative (SFARI) and downloaded the data using Globus Connect Personal v3.2.8.

### Gene Expression Data

Gene expression data in SSC comprised 3,979 samples from probands and siblings across 2,188 families, including 1,791 proband–sibling pairs. Reads were aligned using Spliced Transcripts Alignment to a Reference (STAR version 2.7.9a)^62^ in Weighted Allele-Specific Pipeline (WASP) mode^63^ with reference genome GRCh38. PCR duplicates were marked using Picard MarkDuplicates.^64^ Raw feature counts were computed using the command-line tool featureCounts with duplicates removed and minimum overlap fraction set to 0.5.^65^ All steps up to and including duplicate removal and minimum overlap fraction computation were performed prior to deposition of the data within SFARI.

### Single Nucleotide Polymorphism Data

SNP data included 8,964 samples from 2,358 families. Sequencing was performed on an Illumina HiSeq X10 platform. Reads were aligned to GRCh38 and SNP genotypes were provided by the New York Genome Center (NYGC). Variant discovery was performed using HaplotypeCaller,^66^ followed by Variant Quality Score Recalibration (VQSR)^67^ with parameters maxGaussians = 8 and truth sensitivity = 99.8%. Reads with genotype quality (GQ) < 20 or sequencing depth (DP) < 10 were classified as low quality and removed. All steps up to and including removal of low quality of reads were performed prior to deposition of the data within SFARI.

### Preprocessing

The number of samples retained at each preprocessing step described below is summarized in Supplementary Figure 4.

#### Preprocessing Phenotypic Data

To classify SSC probands as speaking or nonspeaking, we selected individuals assessed using ADOS modules 1–4. These modules correspond to different speech levels: absence of phrase speech, phrase speech, verbal fluency, and verbal fluency in older adolescents. Probands assessed using ADOS Module 1 were classified as nonspeaking, while those assessed with Modules 2–4 were classified as speaking.

Full-scale IQ scores were obtained from the Wechsler Intelligence Scale for Children. Individuals were categorized as follows:

- IQ < 50: moderate-to-profound intellectual disability
- 50 ≤ IQ < 70: mild intellectual disability
- 70 ≤ IQ < 115: no intellectual disability
- IQ ≥ 115: gifted

Samples from two families that withdrew from the study were removed.

#### Preprocessing Gene Expression Data

Gene expression data were imported from SFARI as feature counts. Using the *voom*() function^68^ in the *limma* package (version 3.62.2) for R (version 4.4.3), counts were transformed to log counts per million (*logCPM*) values.^69^ We annotated transcripts using GENCODE basic annotation (version 47).^70^

Analyses included only probands and siblings from each family. Samples lacking relevant phenotypic data were removed. For each gene, we computed a proband–sibling standardized difference by *z*-transforming *logCPM* expression values and subtracting the standardized sibling expression from the standardized proband expression, as defined in Equation 6, where 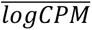 is a vector of *logCPM* mean expression values for each gene and *σ*(*logCPM*) is a vector of *logCPM* standard deviations for each gene.

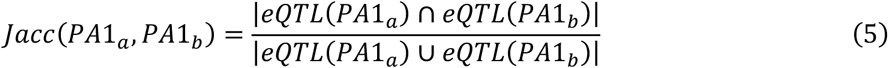

### Differential Genome-Wide Single Nucleotide Polymorphism Associations

Genome-wide association studies (GWAS) were performed while adjusting for ancestry. Using PLINK (version 2.0), we computed SNP-based principal components for all SSC samples (n = 4,240). Fifty principal components were generated after excluding variants with minor allele frequency below 5%. The elbow point on the variance plot was used to determine the number of principal components to retain as ancestry covariates. A covariate file containing ancestry PCs was generated using *awk* (version 4.2.1). Logistic regression GWAS analyses comparing probands with controls were then performed in PLINK 2.0.^71^

### Expression Quantitative Trait Loci

We used PLINK 2.0 to subset the PLINK input files to include only individuals in each PA and ASD subgroup and removed the family ID. We converted the gene expression file to BED format, and mapped genes to the GENCODE Human Genome Release 49 “basic gene annotation” GTF file. We used the Python (version 3.12.12) command-line tool *tensorqtl* ^72^ to compute *p*-values for cis-eQTLs and trans-eQTLs in the PA and ASD subgroups, respectively.

To evaluate overlap between eQTLs in random subsets of PA1 (referred to as *PA*1*_a_* and *PA*1*_b_*), we used the Jaccard index defined in Equation 5. We further computed the mean and standard deviation of the Jaccard index across all random splits.

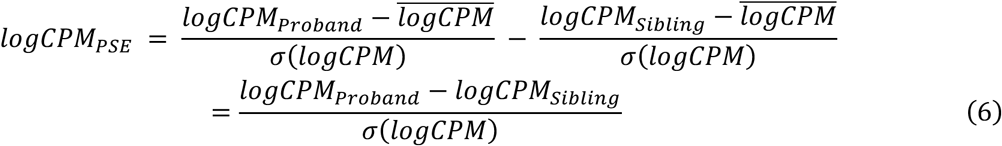

## Data Availability

Gene expression, gene variant (SNP), and phenotypic data used in this analysis are available via controlled access through the Simons Simplex Collection (SSC). The gene sets used by Litman et al. are available through GitHub (https://github.com/FunctionLab/asd-pheno-classes/tree/main/GenomicAnalyses/gene_sets).

## Code Availability

All code used to generate the results presented in this manuscript are available through GitHub (https://github.com/QuackenbushLab/Profound_Autism_Paper_Scripts).

## Acknowledgments

We thank the support staff at the Simons Foundation for assistance with data access. TE and JQ were supported by Amazon Web Services through the Harvard Data Science Institute, and JQ was supported by the National Institutes of Health grant R35CA220523. The authors declare no conflicts of interest.

## Supplementary Tables

Supplementary Table 1. Statistical summary of phenotypic and demographic data for each subgroup.

Supplementary Table 2. Results of Fisher’s Exact Test for each group and each demographic factor.

Supplementary Table 3. Wilcoxon Rank-Sum Test results for *logCPM* in profound and ASD subgroups projected onto each PC (female probands).

Supplementary Table 4. Wilcoxon Rank-Sum Test results for *logCPM* in profound and ASD subgroups projected onto each PC (male probands).

Supplementary Table 5. Wilcoxon Rank-Sum Test results for *logCPM_PSE_* in profound and ASD subgroups projected onto each PC (female probands with male siblings).

Supplementary Table 6. Wilcoxon Rank-Sum Test results for *logCPM_PSE_* in profound and ASD subgroups projected onto each PC (male probands with female siblings).

Supplementary Table 7. Wilcoxon Rank-Sum Test results for *logCPM_PSE_* in profound and ASD subgroups projected onto each PC (female probands with female siblings).

Supplementary Table 8. Wilcoxon Rank-Sum Test results for *logCPM_PSE_* in profound and ASD subgroups projected onto each PC (male probands with male siblings).

Supplementary Table 9. *FDR*-adjusted *p*-values for linear model of *logCPM_PSE_* from PA/ASD subgroup membership.

Supplementary Table 10. *FDR*-adjusted *p*-values for linear model of *logCPM* from PA/ASD subgroup membership.

Supplementary Table 11. FDR-adjusted p-values for Levene’s test of *logCPM* variance equality for PA/ASD subgroups.

Supplementary Table 12. FDR-adjusted p-values for Levene’s test of *logCPM_PSE_* variance equality for PA/ASD subgroups.

Supplementary Table 13. Gene set enrichment analysis results for genes with differential *logCPM* variance between Profound Autism, Nonverbal with IQ ≥ 50 and ASD.

Supplementary Table 14. Gene set enrichment analysis results for genes with differential *logCPM_PSE_* variance between Profound Autism, Nonverbal with IQ ≥ 50 and ASD.

Supplementary Table 15. Mean log-scaled Litman gene set GSEA enrichment scores, including overlapping genes, averaged across 10 iterations.

Supplementary Table 16. Gene sets from Litman et al., formatted as a binary matrix where rows are Ensembl IDs and columns are gene set names. A 1 denotes membership.

Supplementary Table 17. Mean log-scaled Litman gene set GSEA enrichment scores, excluding overlapping genes, averaged across 10 iterations.

Supplementary Table 18. Annotated SNPs associated with autism in at least one subgroup. “chrXY” refers to the pseudo-autosomal region shared by X and Y.

Supplementary Table 19. eQTLs shared across subsets in all ten randomized splits of the PA group meeting both criteria.

