## Supplementary Figures for "Genomic, Transcriptomic, and Regulomic Analyses Do Not Support Profound Autism as a Distinct Biological Category"

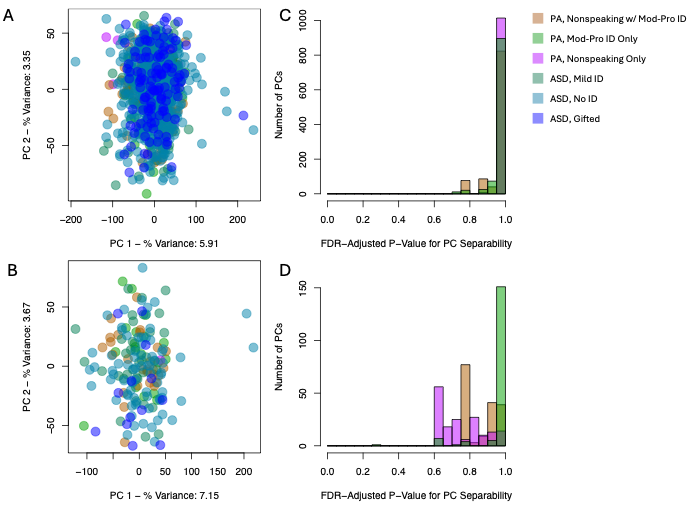

**Supplementary Figure 1. Eigengenes do not exhibit systematic differences between PA and ASD.** Panels (A-B) illustrate each PA and ASD subgroup projected onto the first two eigenvectors, revealing no meaningful subgroup-specific clustering for (A) males and (B) females. Panels (C-D) illustrate the histogram of FDR-adjusted p-values for a Wilcoxon Rank-Sum Test evaluating separability between each PA subgroup and the combined ASD group along each eigengene, revealing no eigengenes that differ significantly between the PA subgroups and the ASD group in (C) males and (D) females.

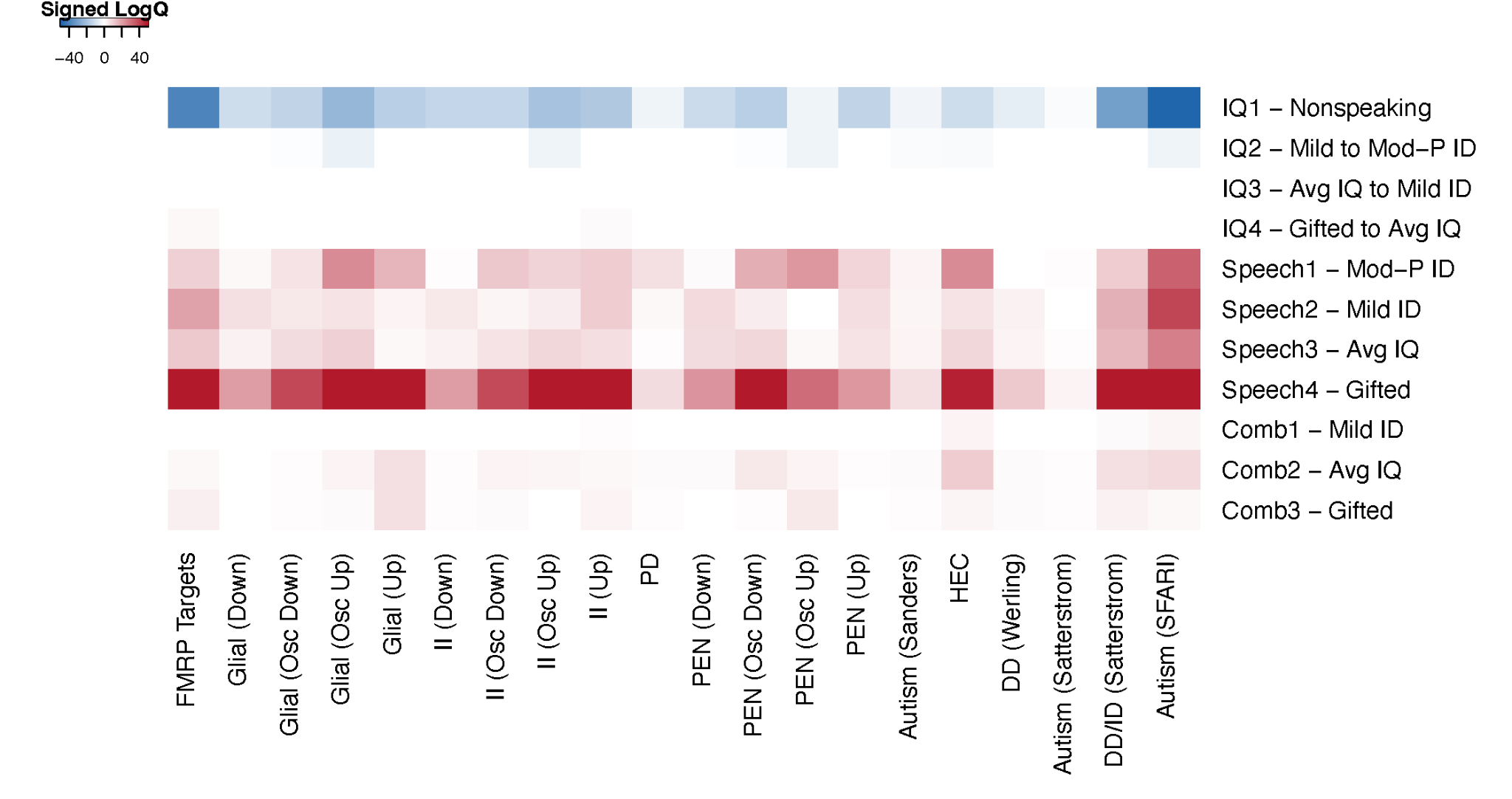

**Supplementary Figure 2. Positive Enrichment Analysis Heatmap for Genes Associated with Autism, Developmental Delay, or Neuronal and Glial Cell Development.** Rows labeled *IQ* refer to comparisons pertaining to decreases in IQ, *Speech* to differences between children with and without phrase speech, and *Comb* to both decreases in IQ and speech capability. Colors correspond to signed log10(*q*-value), with negative enrichment mapped to a negative sign.

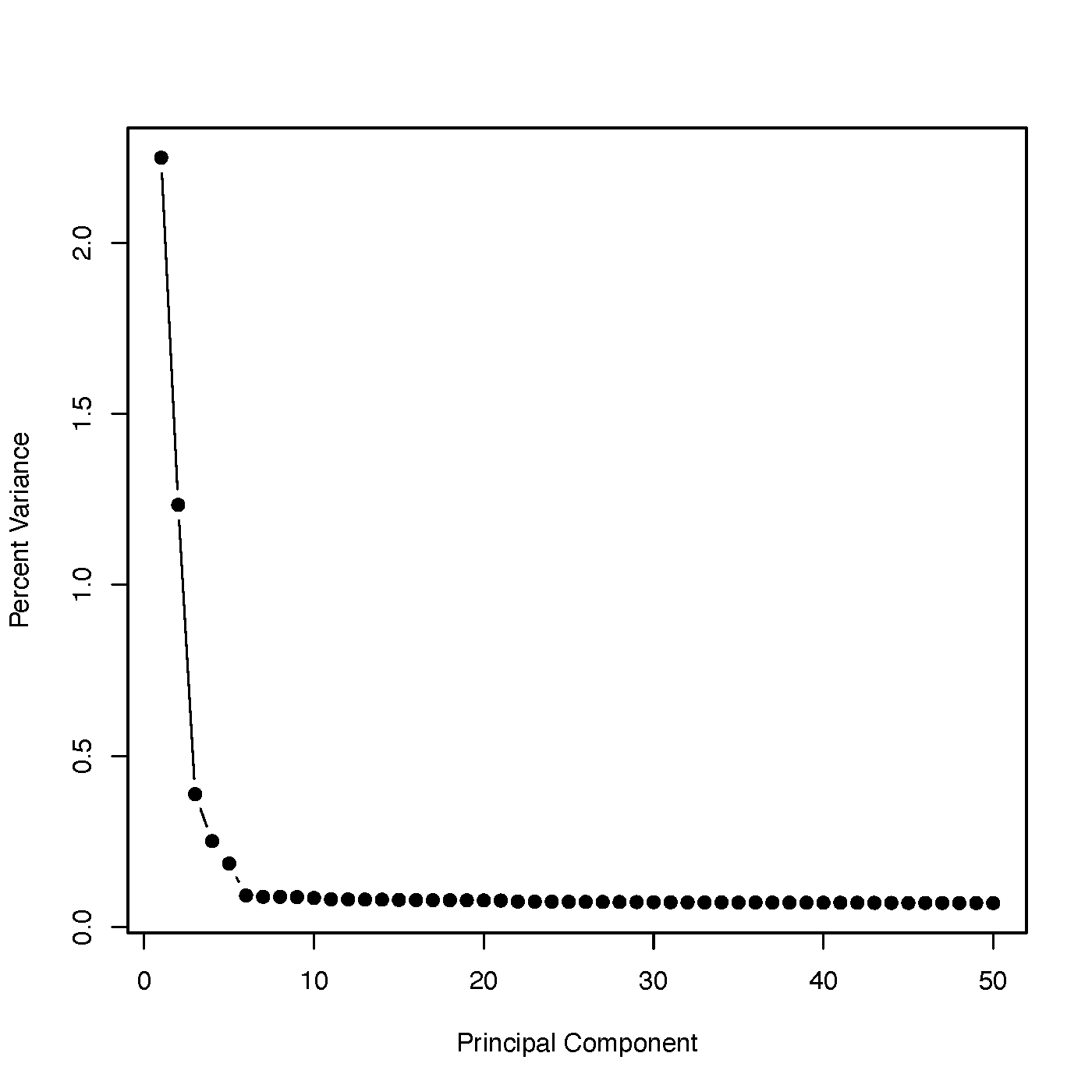

**Supplementary Figure 3. Scree Plot of Percent Variance Explained by SNP-Derived Principal Components.** Principal components were calculated across all samples, including children in the PA and ASD groups and their parents and siblings.

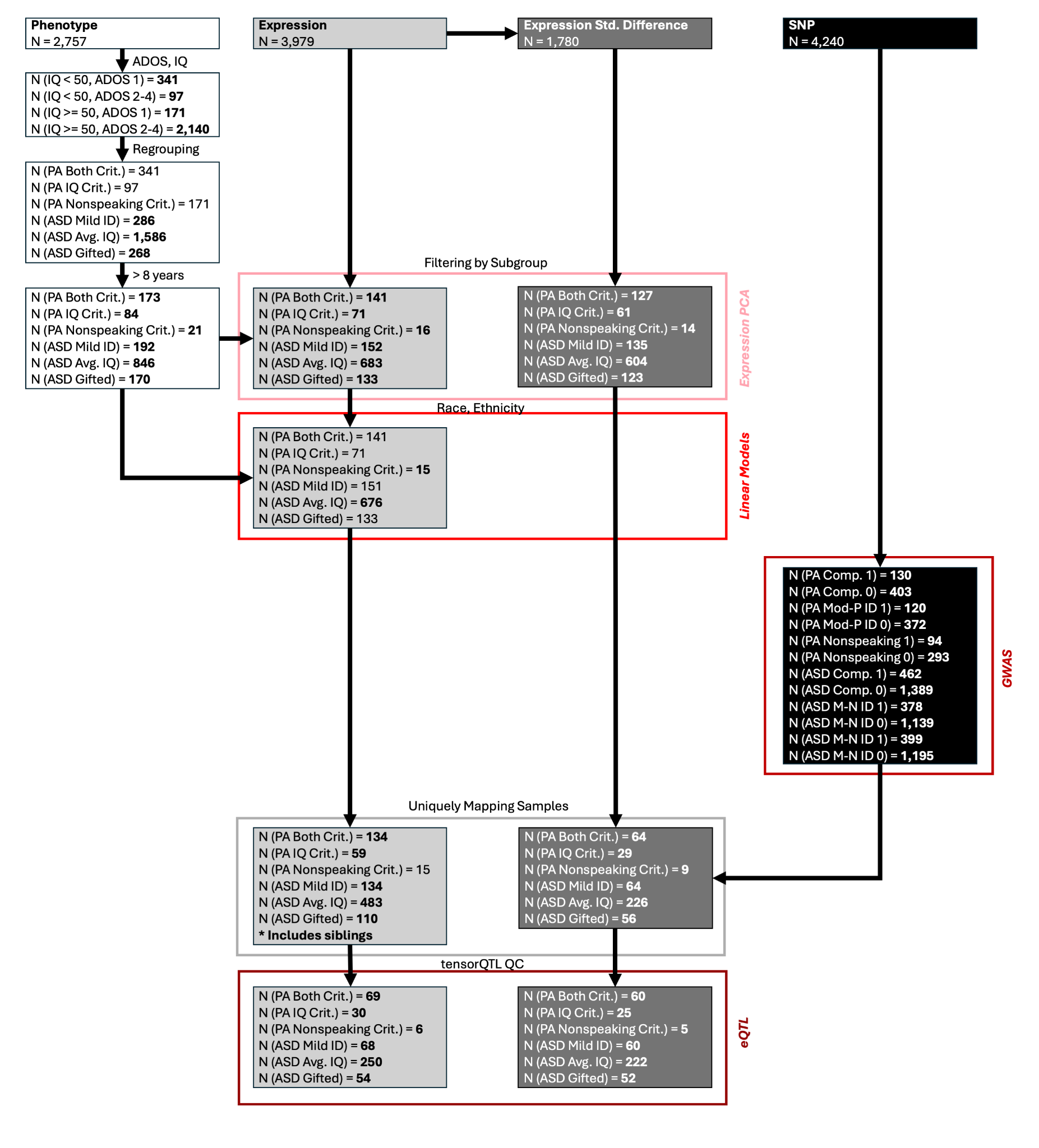

**Supplementary Figure 4. Workflow showing the number of samples retained at each preprocessing step and prior to each analysis.** Each grayscale box refers to a data type used in our analysis, and each arrow to a processing step. Each shade of red corresponds to a different analysis. *N* refers to the number of samples. Bolded values are values that changed after the current processing step.
